# Transfer entropy as a tool for inferring causality from observational studies in epidemiology

**DOI:** 10.1101/149625

**Authors:** N. Ahmad Aziz

## Abstract

Recently Wiener’s causality theorem, which states that one variable could be regarded as the cause of another if the ability to predict the future of the second variable is enhanced by implementing information about the preceding values of the first variable, was linked to information theory through the development of a novel metric called ‘transfer entropy’. Intuitively, transfer entropy can be conceptualized as a model-free measure of directed information flow from one variable to another. In contrast, directionality of information flow is not reflected in traditional measures of association which are completely symmetric by design. Although information theoretic approaches have been applied before in epidemiology, their value for inferring causality from observational studies is still unknown. Therefore, in the present study we use a set of simulation experiments, reflecting the most classical and widely used epidemiological observational study design, to validate the application of transfer entropy in epidemiological research. Moreover, we illustrate the practical applicability of this information theoretic approach to ‘real-world’ epidemiological data by demonstrating that transfer entropy is able to extract the correct direction of information flow from longitudinal data concerning two well-known associations, i.e. that between smoking and lung cancer and that between obesity and diabetes risk. In conclusion, our results provide proof-of-concept that the recently developed transfer entropy method could be a welcome addition to the epidemiological armamentarium, especially to dissect those situations in which there is a well-described association between two variables but no clear-cut inclination as to the directionality of the association.

## Introduction

Epidemiology has been defined as “the study of the distribution and determinants of disease frequency” [1,2]. Therefore, the concept of causation is central to epidemiological research. A uniform definition of causation, however, is lacking and subject to continuing debate and controversy [3-6]. Although several inductively oriented causal criteria are commonly used, especially those proposed by Hill [7], the validity of these criteria for causal inference is regarded as dubious at best [6]. Nevertheless, one criterion of causality which appears to be universally accepted is the condition of temporal precedence: For an exposure to qualify as causal it has to precede the putative effect [6]. Precisely an augmented version of this condition of temporality lies at the heart of the concept of causality as introduced by Norbert Wiener [8]. According to Wiener’s notion of causality one variable could be regarded as the cause of another if the ability to predict the future of the second variable is enhanced by implementing information about the present and past values of the first variable [9]. This concept of causality has been widely applied and validated in various areas of scientific research, most notably in the fields of econometrics and neuroscience [9,10].

Relatively recently Wiener’s concept of causality was linked to information theory, a robust mathematical framework for the quantification of information in the widest sense [11,12]. In the framework of information theory the core measure of information of a variable is its associated Shannon entropy [12,13]. Information entropy represents the reduction of uncertainty which occurs when one actually measures the value of a variable [12-14]. Schreiber demonstrated that if one associates prediction improvement to uncertainty reduction, a measure for Wiener’s causality can be derived within the context of information theory which he defined as transfer entropy [11]. Unlike classical measures of association which are completely symmetric, transfer entropy is an inherently asymmetric measure and is able to detect asymmetries in the interaction of systems evolving in time [11]. Intuitively transfer entropy can be conceptualized as a model-free measure of directed information flow from one variable to another [11]. Therefore, in principle transfer entropy could be a useful measure for deducing causality from observational data in epidemiological research.

Albeit infrequently, information theoretic approaches have been applied before in epidemiology [15-17]. However, their value for inferring causality from observational studies is still unexplored. In the present work, we therefore used a set of simulation experiments as well as real-world data to demonstrate the potential usefulness of the information theoretic metric transfer entropy for causality inference in epidemiological research. Our results provide proof-of-concept that transfer entropy can indeed be applied as a useful tool for assessing causal associations in observational epidemiological studies, especially when there are concerns regarding the direction of action of the presumed causes and effects.

## Methods

### Definitions

The fundamental metric of information theory is Shannon’s entropy which is defined as the average number of bits necessary for optimally encoding a stream of symbols based on the probability of each symbol occurring. Formally, the Shannon’s entropy for variable *X, H*,(*X*) with a probability distribution *p_i_*, is a measure of the expected uncertainty and is given by (the summation is over all states *i* of *X*):

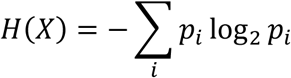

Assuming that two time-series *X* and *Y* could be approximated by Markov processes, then according to Schreiber a measure of causality could be defined as the deviation from the following generalized Markov condition [11,14]:

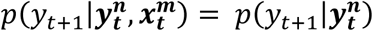

where 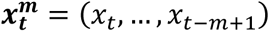 and 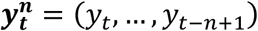 with *m* and *n* the orders of the Markov processes *X* and *Y*, respectively. Now, the deviation from the above Markov condition quantified using the Kullback entropy is defined as the transfer entropy of variable *X* to variable *Y* as follows [11,14]:

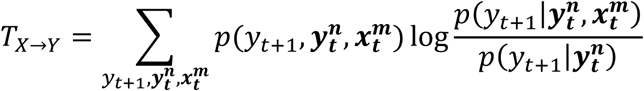

### Simulation experiments

To evaluate the applicability of transfer entropy to observational studies we simulated a series of data sets based on the most fundamental type of epidemiological study, namely a cohort study in which the influence of an exposure *E* is assessed on the occurrence of a disease *D*. For simplicity, the simulations were further based on the following conditions:

1. *N* individuals with a similar age were followed-up for *t* years and data were collected at yearly intervals.
2. Both exposure and disease were assumed to be either present or absent, i.e. both *E* and *D* were represented by a dichotomous variable.
3. The natural rate of occurrence of disease *D* in the unexposed group was assumed to be *p*_0_ per year and the number of disease occurrences was assumed to follow a Poisson distribution.
4. The rate of exposure was assumed to be *p_e_* per year and the number of exposed individuals was assumed to follow a Poisson distribution.
5. There was no loss to follow-up.

A total of four simulated data sets were created based on parameter values for *N, t*, *p_e_* and *p*_0_ of 1000, 10, 0.1 and 0.01, respectively. Each data set was created to represent a different scenario. In scenario **I** we assumed that exposure *E* increased the risk of disease *D* by ten times. In scenario **II** we assumed that disease *D* increased the probability of being exposed by ten times, but that exposure *E* did not increase the risk of disease (i.e. reverse causation). In scenario **III** we assumed that exposure *E* increased the risk of disease *D* by ten times and that disease *D* on its turn also increased the probability of being exposed by ten times. Finally, in scenario **IV** we assumed that a dichotomous confounder *C* increased the probability of being exposed as well as the risk of disease by ten times in the absent of a causal association between *E* and D. In this last scenario, we assumed that the rate of exposure to *C* was 0.1 per year according to a Poisson distribution. For clarity, these four scenarios are represented in **Figure 1** by graphs.

**Figure 1:**
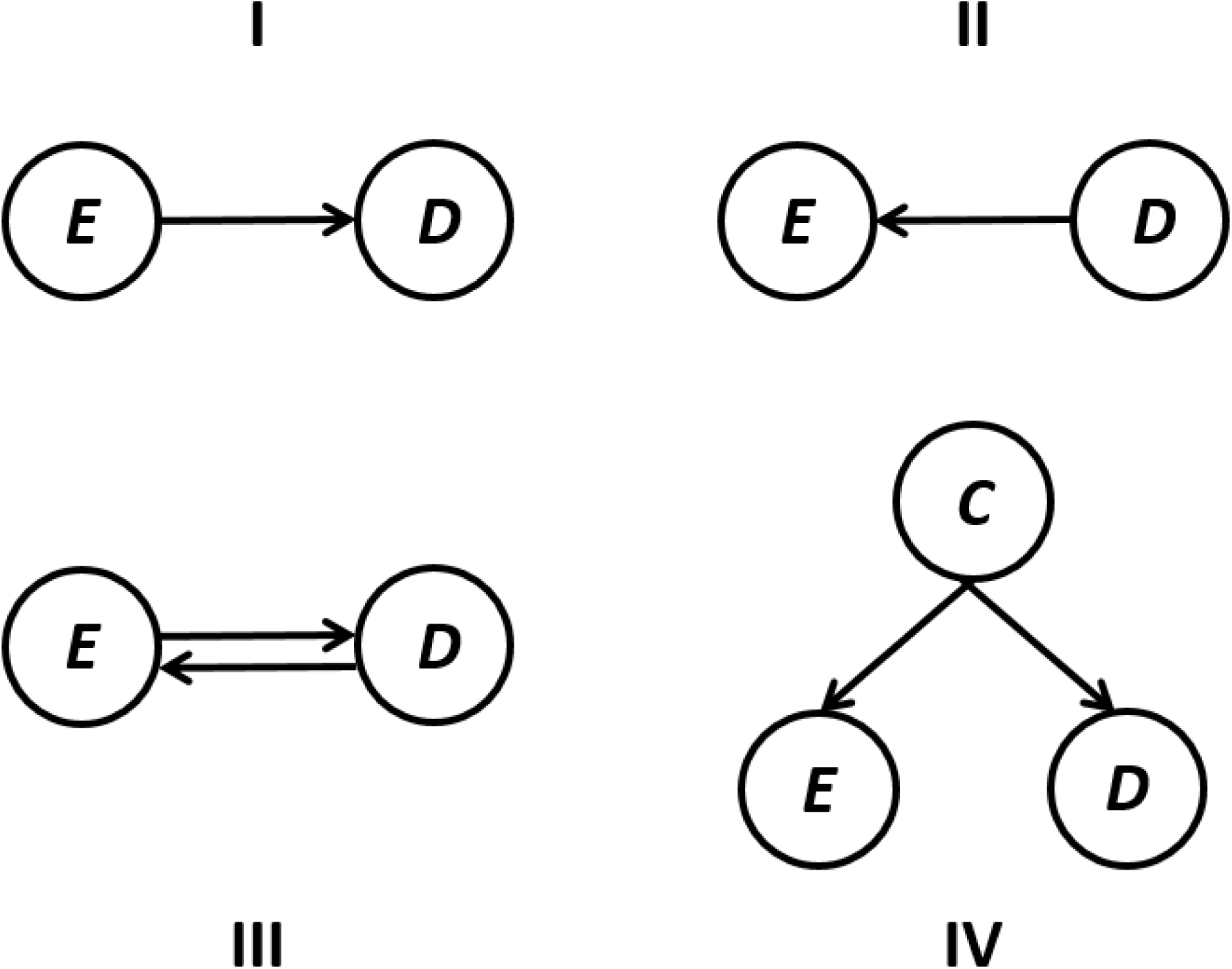
The four simulated scenarios based on the (causal) associations between three different variables (please refer to the text for details).

### Analyses of simulated data

In order to yield robust estimates for transfer entropy each variable had to be converted into a long time-series. This was done for each variable by concatenation of the longitudinal measurements of all simulated subjects, resulting in a long time-series for exposure status and a similar one for disease status. Although by concatenating values for different subjects this method introduces some error into the data, due to the independence of the measurements on each simulated subject this error is completely random and will only result in a conservative estimate, i.e. favoring the null hypothesis of no causal association. For the calculation of the (multivariate) binary transfer entropy metrics we used the Java Information Dynamics Toolkit (version 1.2.1) [18] with the *k* parameter set to follow-up time in order to maximally exploit the information contained in the time-series [11]. Statistical significance for the transfer entropy metrics was determined by computing a distribution of 1000 surrogate values obtained via permutation-based resampling under the null hypothesis of no temporal association between *E* and *D* as described previously [18]. Importantly, for significance testing of multivariate transfer entropy we did not permute the component time-series individually but the vectors of the predictor variables at each time point [19]. Correlations were assessed by the Pearson’s (partial) correlation coefficients while *χ*^2^- or Fisher’s exact tests were applied to assess differences in risk ratios.

### Real-world data

Annual data on age-adjusted lung cancer incidence for all fifty states of the United States of America (USA) were available online for the years 1999 to 2013 through the National Cancer Institute [20]. Data on the percentage of adult daily smokers per state were available online for the period 1995 to 2010 through the Behavioral Risk Factor Surveillance System (BRFSS) [21]. Therefore, we could extract state-level data on both lung cancer incidence and percentage of daily smokers for the period 1999-2010. Similarly, state-wise annual age-adjusted data on the prevalence of obesity (i.e. a body mass index ≥ 30) and diabetes for the years 1995-2015 were available online through the BRFSS [22]. There were only very few missing data values (i.e. < 1%), therefore, missing data points were linearly interpolated.

### Analyses of real-world data

In order to yield robust estimates for transfer entropy we again converted each variable into a long time-series by concatenating the longitudinal measurements of all US states. Given the relatively low number of ‘cases’ (i.e. fifty), the concatenation order was determined completely at random, basing each estimation of transfer entropy on the analysis of 100 random re-orderings of the states. For the calculation of the continuous transfer entropy metrics we used the Kraskov estimator with 4 nearest points using the Java Information Dynamics Toolkit (version 1.2.1) [18]. The history length was set to 1 which in this instance can be interpreted as the extent to which the prevalence of a risk factor (e.g. smoking) could predict the occurrence of a disease (e.g. lung cancer) in the following year. Statistical significance for the transfer entropy for each estimation was again determined by computing a distribution of 1000 surrogate values obtained via permutation-based resampling under the null hypothesis of no temporal association between the two variables of interest [18]. As we also used 100 random re-orderings of the US states for each estimation, the final statistical significance was calculated by bias-corrected bootstrapping of the permutation-based test statistics.

All simulations and analyses were performed using MATLAB Release 2014a (The MathWorks, Inc., Natick, Massachusetts, United States) and SPSS Statistics for Windows Version 21.0 (Armonk, NY: IBM Corp.). All tests were two-tailed and statistical significance level was set at *p* < 0.05.

## Results

### Simulation experiments

#### Scenario I

In this scenario, the correlation coefficient was 0.357 between the two time-series (p <0.001). The transfer entropy of *E* to *D* (*T*_*E*→*D*_) was 0.023 bits and was also highly significant (p <0.001). Conversely, however, the transfer entropy from *D* to *E* (*T*_*D*→*E*_) was only 0.002 bits and was not statistically significant (p = 1.0). Therefore, under this scenario transfer entropy correctly identified the simulated causal relationship in the data which was not apparent from the correlation coefficient alone (**Table 1**).

**Table 1:** Simulated data and analysis of the results at 10 years follow-up.

| *Simulation scenario* |  | *D=0* | *D=1* | *r*<br>(*p-value*)<sup>1</sup> | *partial r*<br>(*p-value*)<sup>1</sup> | *RR*<br>(*p-value*)<sup>2</sup> | *T<sub>E→D</sub>*<br>(*p-value*) | *T<sub>D→E</sub>*<br>(*p-value*) | *T<sub>E→D\|C</sub>*<br>(*p-value*) |
| --- | --- | --- | --- | --- | --- | --- | --- | --- | --- |
| *I* | *E=0* | 345 | 33 | 0.357<br>(<0.001) | - | 4.53<br>(<0.001) | 0.023<br>(<0.001) | 0.002<br>(1.0) | - |
|  | *E=1* | 376 | 246 |  |  |  |  |  |  |
| *II* | *E=0* | 343 | 4 | 0.200<br>(<0.001) | - | 10.50<br>(<0.001) | 0.001<br>(0.359) | 0.010<br>(<0.001) | - |
|  | *E=1* | 574 | 79 |  |  |  |  |  |  |
| *III* | *E=0* | 344 | 8 | 0.321<br>(<0.001) | - | 11.95<br>(<0.001) | 0.010<br>(<0.001) | 0.006<br>(<0.001) | - |
|  | *E=1* | 472 | 176 |  |  |  |  |  |  |
| *IV* | *E=0* | 131 | 17 | 0.271<br>(<0.001) | 0.118<br>(<0.001) | 2.75<br>(*p*<0.001) | 0.006<br>(<0.001) | 0.003<br>(0.990) | 0.004<br>(0.405) |
|  | *E=1* | 583 | 269 |  |  |  |  |  |  |

**Legend:** *D* = disease status (0, no disease; 1, disease); *E* = exposure status (0, no exposure; 1, exposed); *r* = Pearson's correlation coefficient of the two time-series; *RR* = calculated relative risk ratio; *T<sub>E→D</sub>* = transfer entropy from variable *E* to *D* in bits per year; *T<sub>D→E</sub>* = transfer entropy from variable *D* to *E* in bits per year; *T<sub>E→D|C</sub>* = transfer entropy from variable *E* to *D* conditioned on *C* in bits per year
<sup>1</sup>) Assessed by the Pearson's (partial) correlation coefficient between the two time-series.
<sup>2</sup>) Statistical significance was assessed by the $\chi^2$ -test except when cell counts were smaller than 10, in which case the Fisher's exact test was applied.

#### Scenario II

In this scenario of ‘reverse causation’ both the correlation coefficient and the risk ratio as calculated from a contingency table were highly significant (**Table 1**). However, only the transfer entropy metrics could identify the true direction of the effect, namely that the disease increased the risk of exposure and not vice versa, reflected in a statistically significant *T*_*D*→*E*_ but not *T*_*E*→*D*_ (**Table 1**).

#### Scenario III

In this scenario of mutual influence a statistically highly significant association between exposure and disease was indeed detected by the correlation coefficient and risk ratio metrics. Again, however, the bilateral direction of this ‘causation’ only became apparent through transfer entropy analysis: both *T*_*E*→*D*_ and *T*_*D*→*E*_ were statistically significant (**Table 1**).

#### Scenario IV

In this scenario, a spurious association was created between the exposure and disease through a confounding variable *C* which strongly influenced both *E* and *D* (**Figure 1**). As expected all univariate metrics detected an association between *E* and *D*. The difference between the metrics became only apparent when we corrected for the effect of variable *C*. Although the partial correlation between *E* and *D* (adjusted for *C*) was less than half of the uncorrected correlation, it still remained highly significant (**Table 1**). Conversely, the conditional entropy between *E* and *D* – which was calculated using the formula *T*_*E*→*D*|*C*_ = *T*_*E*,*C*→*D*_ − *T*_*C*→*D*_ (see the **appendix** for a formal proof of this identity) – was not statistically significant (**Table 1**), thus correctly eliminating the spurious association mediated through the confounder *C*.

### Real-world data

#### Smoking and lung cancer incidence

As expected there was a highly significant correlation between the two time-series representing the percentage of daily smokers and the incidence of lung cancer (bootstrapped values: r = 0.719, p << 0.001; **Figure 2**). Interestingly, the transfer entropy metrics in addition could identify the directionality of the effect: The transfer entropy from smoking to lung cancer incidence (i.e. *T*_*Smoking*→*Lung cancer*_) was highly significant (bootstrapped values: 0.058 bits, p = 0.008), whereas the transfer entropy from lung cancer incidence to smoking (i.e. *T*_*Lung cancer*→*Smoking*_) was not (bootstrapped values: 0.005 bits, p = 0.813).

**Figure 2:**
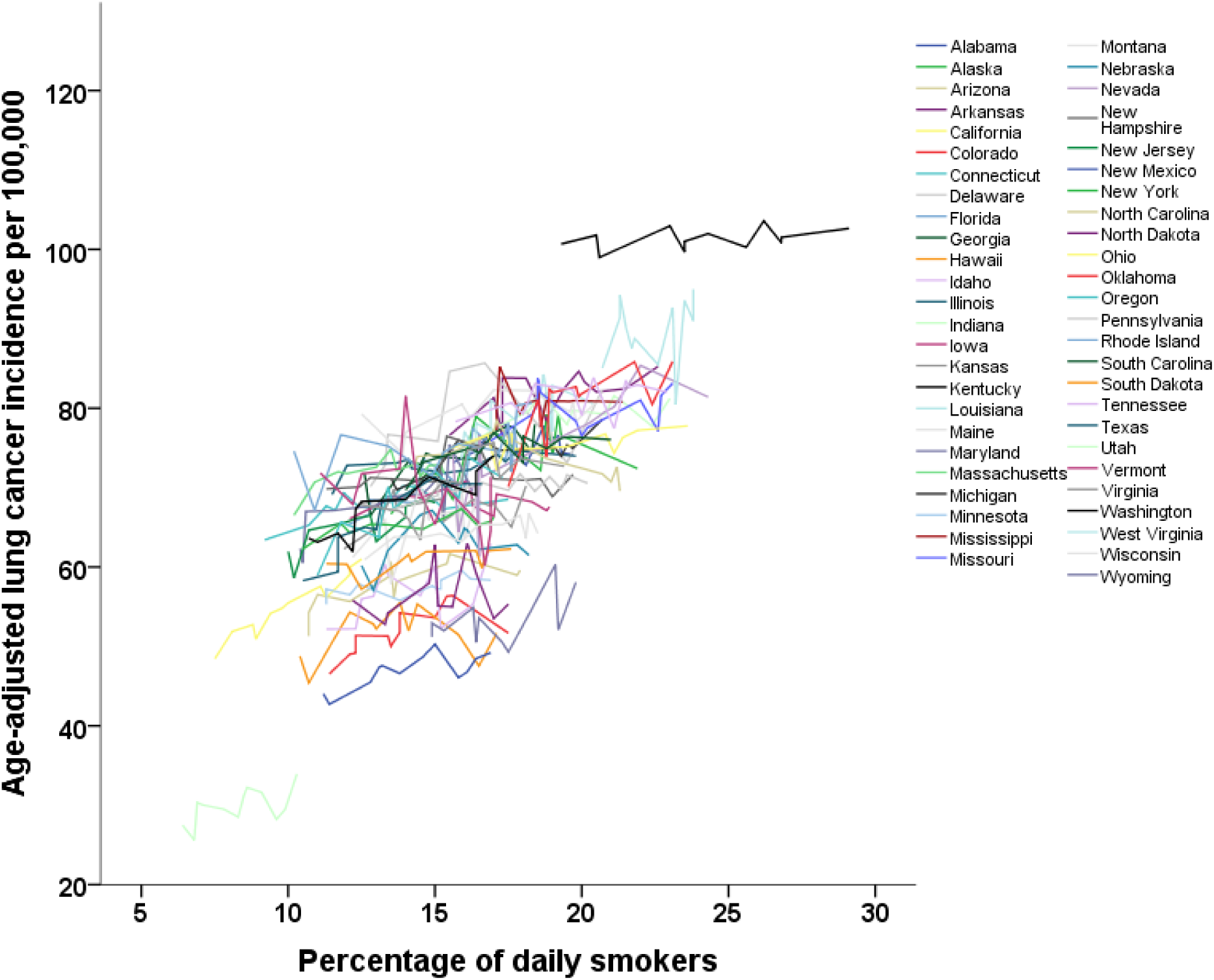
The incidence of lung cancer during the period 1999-2010 is strongly associated with the percentage of daily smokers in the USA. Each line represents a different US state.

#### Obesity and diabetes prevalence

The two time-series containing the prevalence of obesity and diabetes were highly correlated as expected (bootstrapped values: r = 0.888, p << 0.001, **Figure 3**). Again, transfer entropy analysis was able to distinguish between the putative cause and effect: The transfer entropy from obesity prevalence to diabetes prevalence (i.e. *T*_*Obesity*→*Diabetes*_) was highly significant (bootstrapped values: 0.131 bits, p << 0.001), whereas the transfer entropy from diabetes prevalence to obesity prevalence (i.e. *T*_*Diabetes*→*Obesity*_) was not (bootstrapped values: 0.054 bits, p = 0.236).

**Figure 3:**
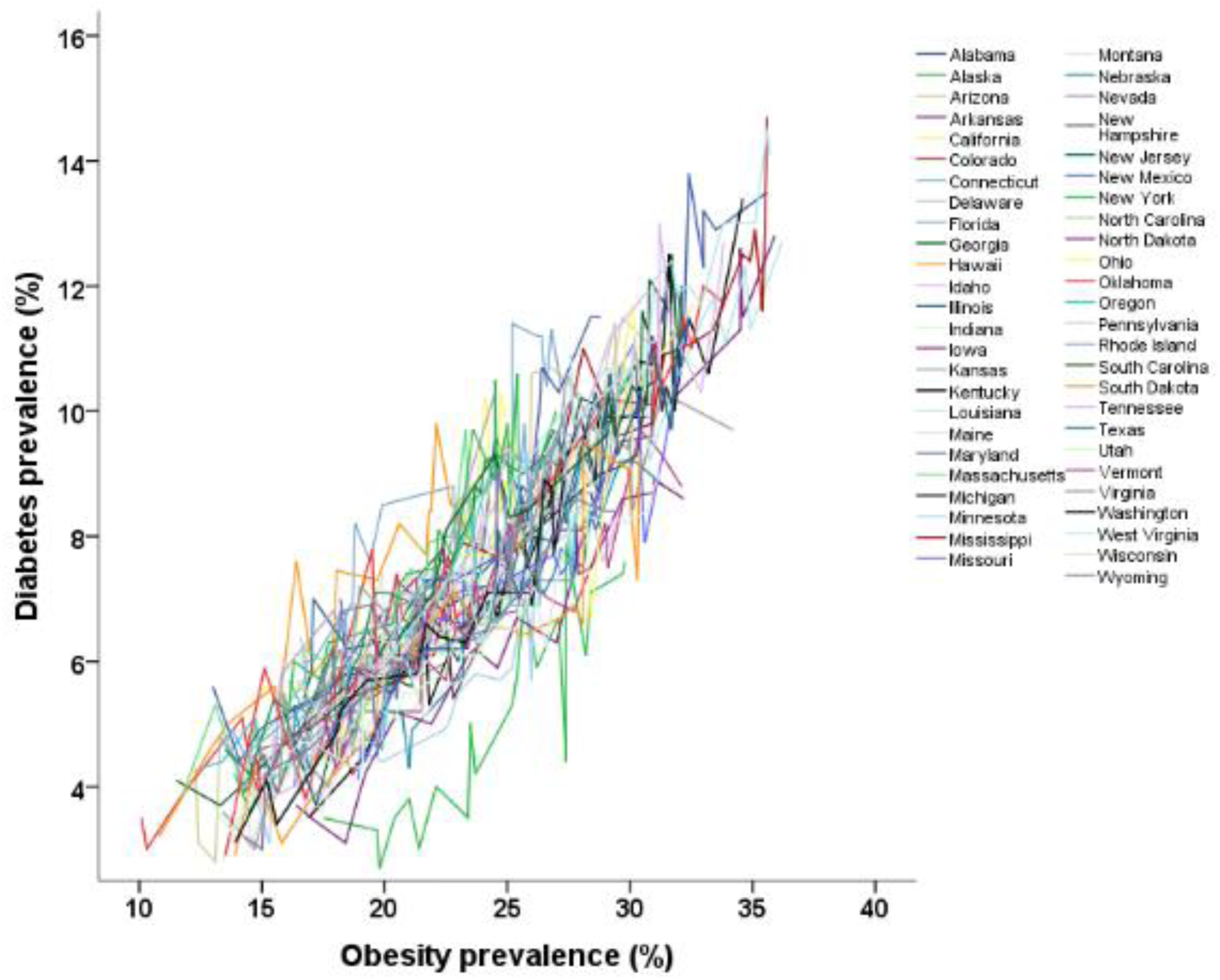
The prevalence of obesity and diabetes during the period 1995-2015 are strongly associated in the USA. Each line represents a different US state.

## Discussion

One of the greatest challenges of epidemiological research is to infer causal relationships from non-experimental observations [6]. In epidemiology, this challenge is in fact reflected in a fundamental dichotomy between ‘observational’ and ‘experimental’ studies because only the latter are deemed to be able to provide unequivocal evidence for or against a causal effect [23,24]. Epidemiologists however do not stand alone in facing this grand obstacle since the issue of inferring causality from non-experimental data is pervasive to all disciplines of science concerned with etiological research: From the neurosciences in which massive amounts of connections between various parts of the brain have to be evaluated for ‘effective connectivity’, i.e. identifying the ‘driver’ and ‘responder’ components of neural networks [19,25], to cause-effect relationships in various climate models [26], to detecting causal links between different financial markets [10,27], just to name a few examples. The recent merging of Wiener’s causality theorem with information theory and the resulting notion of information flow as transfer entropy has revolutionized research in all these and many other disciplines [11,14], but surprisingly, not in epidemiology.

In this paper, we therefore aimed to introduce the concept of transfer entropy and its potential applicability to epidemiological research. In order to assess the validity of transfer entropy when applied to epidemiological data we specifically simulated a series of data sets to obtain ground truth about the underlying causal associations which obviously would have not been possible in a ‘real’ data set. The simulations we chose concerned the most classical, basic and widely used epidemiological observational study design: the assessment of the effects of a dichotomous exposure on a dichotomous outcome in a cohort of individuals who were followed up for a certain amount of time [1,2]. Our results show that transfer entropy can indeed be used as a powerful tool to assess causal associations in observational epidemiological studies. Importantly, comparison with traditional epidemiological metrics showed that the application of transfer entropy can provide valuable insight into the direction of information flow. Directionality of information flow is simply not reflected in traditional (linear) measures of association which are completely symmetric by design, nicely reflected in the old adage “association is not causation”. This symmetric design of traditional measures of association implies that for the evaluation of the direction of association additional information will be necessary, whereas transfer entropy can deduce the direction of information flow from the data at hand [11].

In order to illustrate the practical applicability of this information theoretic approach to ‘real-world’ epidemiological data, we also assessed whether transfer entropy was able to extract the correct direction of information flow from longitudinal data on two well-known associations. The first association concerned that between smoking and lung cancer, while the second relation concerned that between obesity and diabetes risk [28,29]. We chose these examples as in both instances there is ample theoretical and experimental evidence to consider one of the variables as the cause and the other as the effect, i.e. smoking and obesity cause lung cancer and diabetes, respectively, and not vice versa [28,29]. In both examples, our analysis demonstrated that transfer entropy could correctly pinpoint the direction of information flow, and thereby discern the cause from the effect, lending further support to the validity of this approach.

Our results also demonstrate another notable advantage of transfer entropy over traditional measures of association, namely the fact that transfer entropy is a model-free measure of information flow [11]. An important corollary thereof is that in case no information is available about the underlying data structure, and especially when non-linear interactions are possible, transfer entropy is more likely to yield valid results as compared to model-based approaches. In our simulated data sets this advantage was most evident in scenario IV in which we had created a spurious association between an exposure and a disease completely mediated via a confounder which influenced both. In this scenario adjusting the association between exposure and disease for the confounder through application of partial correlation, which assumes linearity, only partly removed the confounding effect. In contrast, the transfer entropy of the association between the exposure and disease indicated no information flow when adjusted for the influence of the confounder. Although, obviously, stratification by the confounder would also have eliminated the spurious relationship between exposure and disease in this simple example with a dichotomous confounding variable, this example does illustrate that linear correction does not necessarily remove all confounding effects. Moreover, in more complex cases stratification becomes problematic and statistical adjustment will be required. The availability of multivariate transfer entropy methods is therefore very useful in cases were intricate association networks have to be assessed, a situation which is for example very common in the neurosciences where transfer entropy has proven to be very useful in the assessment of dozens of pathways in order to establish ‘effective connectivity’ [19]. Furthermore, it should be noted that recently it was proven that transfer entropy reduces to Granger causality, a linear measure of causality, for normally distributed data [30,31].

It is also important to note that while in our simulations all variables were assumed to be dichotomous for simplicity, there is no reason to restrict the application of transfer entropy in epidemiology to binary data as exemplified by our ‘real-world’ data analyses. In fact there are very well described estimation methods for transfer entropy for both discrete as well as continuous data [18,32]. Additionally, these methods are readily available as they are conveniently implemented in various freely available software toolboxes and scripts for use in various programming languages [18,32-34]. Lizier provides a comprehensive overview and discussion of the currently available software [18].

Despite the advantages of transfer entropy as a measure of directed association there are also some limitations to this method [14]. First, as transfer entropy is based on Wiener’s principle of causality which pertains to non-interventional, observational causality, it is like all traditional methods liable to the effects of unobserved confounders. The availability of multivariate transfer entropy does however allow for conditioning on the effects of observed variables which could potentially act as confounders. Secondly, the fact that transfer entropy is a model-free measure of information flow automatically entails that although it can accurately detect the direction and magnitude of information flow, it cannot describe the nature of the information conveyed. However, this problem can be overcome by using post-hoc model-based approaches to assess the specific nature of the interaction after significant information flow pathways have been identified through application of transfer entropy [14]. Another consequence of a model-free measure is that there is no fixed probability distribution, although statistical significance testing is possible through resampling methods as described here.

In conclusion, in this study we provide proof-of-concept for the usefulness of transfer entropy as a measure of directed association in observational studies in epidemiology. Our findings indicate that the recently developed transfer entropy method could be a welcome addition to the epidemiological armamentarium, especially to dissect those situations in which there is a well-described association between two variables but no clear-cut inclination as to the directionality of the association. Examples of the latter include the association between atrial fibrillation and cryptogenic stroke [35], the relation between statin use and cognitive decline [36] and the putative link between cancer and diabetes [37], although there are myriads more of such clinically relevant associations which could be subjected to further scrutiny by transfer entropy analysis. Another potential use could be in observational follow-up studies in which large amounts of variables are periodically measured; in the analysis of data from these studies transfer entropy could be applied to reconstruct the contours of the information flow in an unbiased manner analogous to reconstruction of networks of effective connectivity from functional imaging data [19].

## Appendix

The transfer entropy of a stochastic time-varying variable *X* on another *Y* can also be expressed in terms of their associated conditional Shannon entropies[32]:

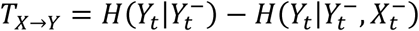

Where the subscript *t* represents the present time, and 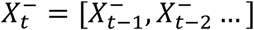 and 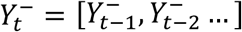 denote vectors representing the whole past of the variables *X* and *Y*. Similarly, the multivariate transfer entropy of two variables *X* and *Z* on *Y* is given by[32]:

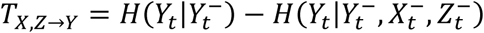

Where 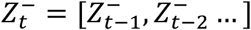 denotes a vector representing the whole past of the variable *Z*. From the above it follows immediately that:

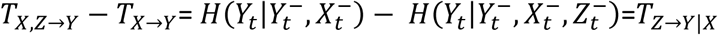

With *T*_*Z*→*Y*|*X*_ thus representing the transfer entropy from variable *Z* to *Y* conditioned on *X*.

## Acknowledgements

The author would like to thank Prof. dr. S. le Cessie for comments on an earlier version of this manuscript.

## Supporting data

Matlab scipts
Dataset 1: Smoking & Lung cancer (1999-2010)
Dataset 2: Obesity & Diabetes prevalence (1995-2015)

## References

1. MacMahon B, Pugh TF (1970) Epidemiology: Principles and Methods. Boston: Little, Brown.

2. Rothman KJ (2012) Epidemiology: An Introduction. New York: Oxford University Press, Inc.

3. Parascandola M, Weed DL (2001) Causation in epidemiology. J Epidemiol Community Health 55: 905-912.

4. Olsen J (2003) What characterises a useful concept of causation in epidemiology? J Epidemiol Community Health 57: 86-88.

5. Pearl J (2010) An introduction to causal inference. Int J Biostat 6: Article.

6. Rothman KJ, Greenland S (2005) Causation and causal inference in epidemiology. Am J Public Health 95 Suppl 1: S144-S150.

7. Hill AB (2015) The environment and disease: association or causation? J R Soc Med 108: 32-37.

8. Wiener N (1956) The theory of prediction. Modern mathematics for engineers. New York: McGraw-Hill.

9. Bressler SL, Seth AK (2011) Wiener-Granger causality: a well established methodology. Neuroimage 58: 323-329.

10. Granger CWJ (1969) Investigating Causal Relations by Econometric Models and Cross-spectral Methods. Econometrica 37: 424-438.

11. Schreiber T (2000) Measuring information transfer. Phys Rev Lett 85: 461-464.

12. Shannon C (1948) A mathematical theory of communication. Bell System Technical Journal 27: 379-423.

13. Reza FM (2012) An introduction to information theory. New York: Dover Publications, Inc.

14. Vicente R, Wibral M, Lindner M, Pipa G (2011) Transfer entropy--a model-free measure of effective connectivity for the neurosciences. J Comput Neurosci 30: 45-67.

15. Mandava P, Krumpelman CS, Shah JN, White DL, Kent TA (2013) Quantification of errors in ordinal outcome scales using shannon entropy: effect on sample size calculations. PLoSONE 8: e67754.

16. Sumi A, Kamo K, Ohtomo N, Mise K, Kobayashi N (2011) Time series analysis of incidence data of influenza in Japan. J Epidemiol 21: 21-29.

17. Rhodes CJ, Demetrius L (2010) Evolutionary entropy determines invasion success in emergent epidemics. PLoSONE 5: e12951.

18. Lizier JT (2014) JIDT: An information-theoretic toolkit for studying the dynamics of complex systems. Frontiers in Robotics and AI 1.

19. Lizier JT, Heinzle J, Horstmann A, Haynes JD, Prokopenko M (2011) Multivariate information-theoretic measures reveal directed information structure and task relevant changes in fMRI connectivity. J Comput Neurosci 30: 85-107.

20. U.S. Cancer Statistics Working Group. United States Cancer Statistics: 1999–2013 Incidence and Mortality Web-based Report. Available at: http://www.cdc.gov/uscs. Atlanta (GA): Department of Health and Human Services, Centers for Disease Control and Prevention, and National Cancer Institute.

21. Centers for Disease Control and Prevention (CDC). Behavioral Risk Factor Surveillance System Survey Data. Available at: https://chronicdata.cdc.gov/Survey-Data/Behavioral-Risk-Factor-Data-Tobacco-Use-2010-And-P/fpp2-pp25. Atlanta, Georgia: U.S. Department of Health and Human Services, Centers for Disease Control and Prevention.

22. Centers for Disease Control and Prevention (CDC). Behavioral Risk Factor Surveillance System Survey Data. Available at: https://chronicdata.cdc.gov/Survey-Data/Behavioral-Risk-Factor-Data-Tobacco-Use-2010-And-P/fpp2-pp25. Atlanta, Georgia: U.S. Department of Health and Human Services, Centers for Disease Control and Prevention.

23. Vandenbroucke JP (2009) The HRT controversy: observational studies and RCTs fall in line. Lancet 373: 1233-1235.

24. Vandenbroucke JP (2004) When are observational studies as credible as randomised trials? Lancet 363: 1728-1731.

25. Wibral M, Rahm B, Rieder M, Lindner M, Vicente R, et al. (2011) Transfer entropy in magnetoencephalographic data: quantifying information flow in cortical and cerebellar networks. ProgBiophysMolBiol 105: 80-97.

26. Runge J, Heitzig J, Petoukhov V, Kurths J (2012) Escaping the curse of dimensionality in estimating multivariate transfer entropy. Phys Rev Lett 108: 258701.

27. Dimpfl T, Peter FJ (2013) Using transfer entropy to measure information flows between financial markets. Studies in Nonlinear Dynamics and Econometrics 17: 85-102.

28. Lee PN, Forey BA, Coombs KJ (2012) Systematic review with meta-analysis of the epidemiological evidence in the 1900s relating smoking to lung cancer. BMC Cancer 12: 385.

29. Eckel RH, Kahn SE, Ferrannini E, Goldfine AB, Nathan DM, et al. (2011) Obesity and type 2 diabetes: what can be unified and what needs to be individualized? Diabetes Care 34: 1424-1430.

30. Barnett L, Barrett AB, Seth AK (2009) Granger causality and transfer entropy are equivalent for Gaussian variables. Phys Rev Lett 103: 238701.

31. Hlavácková-Schindler K (2011) Equivalence of granger causality and transfer entropy: A generalization. App Math Sci 5: 3637-3648.

32. Montalto A, Faes L, Marinazzo D (2014) MuTE: a MATLAB toolbox to compare established and novel estimators of the multivariate transfer entropy. PLoSONE 9: e109462.

33. Astakhov S, Grassberger P, Kraskov A, Stögbauer H (2013) Mutual information least-dependent component analysis (MILCA). Available from: http://wwwuclacuk/ion/departments/sobell/Research/RLemon/MILCA/MILCA.

34. Niso G, Bruna R, Pereda E, Gutierrez R, Bajo R, et al. (2013) HERMES: towards an integrated toolbox to characterize functional and effective brain connectivity. Neuroinformatics 11: 405-434.

35. Favilla CG, Ingala E, Jara J, Fessler E, Cucchiara B, et al. (2015) Predictors of Finding Occult Atrial Fibrillation After Cryptogenic Stroke. Stroke.

36. Power MC, Weuve J, Sharrett AR, Blacker D, Gottesman RF (2015) Statins, cognition, and dementia-systematic review and methodological commentary. NatRevNeurol 11: 220-229.

37. Harding JL, Shaw JE, Peeters A, Cartensen B, Magliano DJ (2015) Cancer risk among people with type 1 and type 2 diabetes: disentangling true associations, detection bias, and reverse causation. Diabetes Care 38: 264-270.

